# Osterix-driven LINC complex disruption *in vivo* diminishes bone microarchitecture in 8-week male mice but not after 6-week voluntary wheel running

**DOI:** 10.1101/2023.08.24.554623

**Authors:** Scott Birks, Sean Howard, Caroline O’Rourke, William R Thompson, Anthony Lau, Gunes Uzer

**Affiliations:** Boise State University, Micron School of Materials Science and Engineering; Boise State University, Mechanical and Biomedical Engineering; The College of New Jersey, Biomedical Engineering; Indiana University, Department of Physical Therapy

**Keywords:** Osterix, Mechanical Signals, Mechanobiology, LINC, Bone

## Abstract

The Linker of Nucleoskeleton and Cytoskeleton (LINC) complex is a crucial connective component between the nuclear envelope and the cytoskeleton involving various cellular processes including nuclear positioning, nuclear architecture, and mechanotransduction. How LINC complexes regulate bone formation *in vivo*, however, is not well understood. To start bridging this gap, here we created a LINC disruption murine model using transgenic mice expressing Cre recombinase enzyme under the control of the Osterix (Osx-Cre) which is primarily active in pre-osteoblasts and floxed Tg(CAG-LacZ/EGFP-KASH2) mice. Tg(CAG-LacZ/EGFP-KASH2) mice contain a lox-STOP-lox flanked LacZ gene which is deleted upon cre recombination allowing for the overexpression of an EGFP-KASH2 fusion protein. This overexpressed protein disrupts endogenous Nesprin-Sun binding leading to disruption of LINC complexes. Thus, crossing these two lines results in a <u>O</u>sx-<u>d</u>riven <u>L</u>INC <u>d</u>isruption (ODLD) specific to pre-osteoblasts. In this study, we investigated how this LINC disruption affects exercise induced bone accrual. ODLD cells had decreased osteogenic and adipogenic potential *in vitro* compared to non-disrupted controls and sedentary ODLD mice showed decreased bone quality at 8-weeks. Upon access to a voluntary running wheel ODLD animals showed increased running time and distance; however, our 6-week exercise intervention did not significantly affect bone microarchitecture and bone mechanical properties.

## Introduction

The ability of bone progenitor cells to sense and respond to mechanical signals is fundamental to bone homeostasis and remodeling mechanisms [1]. A crucial player in cellular mechanotransduction pathways is the Linker of Nucleoskeleton and Cytoskeleton (LINC) complex, which serves as a physical link between the cytoskeleton and the nucleoskeleton, facilitating the transmission of mechanical forces and signals between the extracellular matrix and the nucleus [2].

LINC complexes are protein assemblies that connect the cytoskeleton to the nucleus through interactions between cytoskeletal nesprins (nuclear envelope spectrin repeats) and perinuclear Sun (Sad1/Unc84) proteins. The N-terminal calponin homology domains of nesprins bind to cytoskeletal F-actin microfilaments, while the evolutionarily conserved C-terminal KASH (Klarsicht/Anc-1, Syne Homology) domains bind to Sun domains within the perinuclear space. Sun proteins directly engage with the intranuclear lamina, completing the connection from the cytoskeleton to the nucleus [3–5]. Beyond their structural role, LINC complexes have emerged as critical components in transducing mechanical signals within cells[6].

Previous *in vitro* studies have demonstrated the active role of LINC complexes in mechanotransduction pathways [7, 8]. For example, via inhibiting the LINC complex using RNA interference to suppress SUN1 and SUN2 proteins or transfecting cells with a dominant negative form of the Nesprin KASH domain results in the muting of FAK and Akt phosphorylation and inhibit the mechanical repression of adipogenesis when mesenchymal stromal cells are exposed to low-magnitude, high-frequency signals [7]. While the importance of LINC complexes in mechanotransduction has been evidenced *in vitro*, their role in mediating mechanical regulation of bone *in vivo* remains largely unexplored. The present study builds upon these cellular insights by employing an *in vivo* approach to examine the effects of LINC complex disruption under mechanical loading specifically in pre-osteoblastic cells using an Osterix-driven genetic model.

Osterix (Osx or Sp7) is a transcription factor indispensable for osteoblastogenesis and subsequent bone formation[9, 10]; therefore, the Osx-Cre driver was selected to enable the investigation of LINC complex disruption effects in cells already committed to the osteogenic lineage [11]. LINC disruption was achieved using floxed mice created by Razafsky and Hodzic, which overexpresses an EGFP-KASH2 fusion protein upon Cre-recombination [12, 13]. This fusion protein binds to perinuclear Sun proteins, causing displacement of cytoplasmic nesprins into the endoplasmic reticulum and thus, disrupting LINC connectivity between the nucleus and cytoskeleton.

Exercise is considered a potent modality of mechanical stimulation that can trigger adaptive responses in bone tissue, enhancing bone density and strength [14]. Multiple rodent exercise models exist including forced/trained treadmill running, voluntary/forced wheel running, swimming, and resistance training (loaded/unloaded ladder climbing), all of which have been shown to improve bone mass, bone strength, and bone mineral density [15] [16]. In this study, we have chosen voluntary wheel running model as it is less stressful to the animal.

We hypothesized that disabling LINC function decreases osteogenesis and will hinder exercise-induced bone accrual *in vivo*. To interrogate this question, we created a LINC disruption model under the Osx-Cre driver and measured primary cell differentiation potential *in vitro*, bone microarchitecture via mico-computerized tomography (Micro-CT) and biomechanical properties via 3-point bending at an 8-week baseline. We then subjected a cohort of animals to either a running or non-running intervention starting at 8-weeks of age and lasting for 6 weeks with a 1-week acclimation period. Micro-CT and biomechanical properties were also measured post-exercise to assess any effects of LINC disruption on exercise induced bone accrual.

## Materials and Methods

### Animals

Mouse strains included B6.Cg-Tg(Sp7-tTA,tetO-EGFP/cre)1Amc/J aka *Osx-Cre* (The Jackson Laboratory, Stock Number: 006361)[11], Tg(CAG-LacZ/EGFP-KASH2) aka *KASH2* (Hodzic, Wash U)[12], and B6.Cg-Gt(ROSA)26Sor^tm14(CAG-tdTomato)Hze^/J aka *Ai14* (generously provided by Dr. Richard Beard)[17].

Hemizygous *Osx-Cre* mice were crossed with floxed *KASH2* mice to generate an osterix-driven LINC disrupted (ODLD) murine model. LINC disruption mechanism described by Razafsky and Hodzic [12, 13]. Briefly, upon cre recombination, a LacZ containing lox-stop-lox cassette upstream of an eGFP-KASH2 ORF is excised allowing for the overexpression an eGFP-KASH2 fusion protein which saturates available Sun/KASH binding in a dominant-negative manner. Genotyping was performed on tissue biopsies via real-time PCR probing for LacZ and Cre (Transnetyx, Cordova, TN) to determine experimental animals and controls. Cre(+)/LacZ(+) animals were considered experimental and Cre(+)/LacZ(-) mice were used as controls due to known skeletal phenotype associated with Osx-Cre(+) animals. Mice were not treated with tetracycline or its derivatives as Osx-Cre activation did not cause significant fetal lethality. Animals were housed in individually ventilated cages under controlled conditions until placement in conventional cages for the duration of the exercise intervention. *Ad libitum* access to food and water allowed.

Additionally, *Osx-Cre* mice were crossed with Ai14 RFP reporter mice to assess Osx-Cre activity in skeletal tissue and bone marrow cells. All procedures approved by Boise State University Institutional Animal Care and Use Committee.

### Bone Marrow Cell Isolation, Expansion, and Differentiation

Bone marrow cells were isolated from tibiae and femora via centrifugation as previously described [18]. Briefly, tibia and femur were dissected, cleared of all soft tissue, and centrifuged at >12,000g to remove the bone marrow. This bone marrow pellet was then resuspended in media and plated in supplemented growth media (alpha-MEM +20% FBS, +100U/mL penicillin, +100µg/mL streptomycin). Cells were washed with PBS after 48 hours to remove non-adherent cells and allowed to proliferate to 50% confluency. At this confluency, cells were lifted, split onto two separate plates to produce enough cells for assays, and allowed to proliferate to ∼80% confluency. Cells were then lifted and plated for differentiation assays.

For osteogenic assays (n=2 mice /group), cells were seeded at 200K per well on 6-well plates in normal growth media for 24 hours prior to switching to osteogenic media (alpha-MEM with 20% FBS, 100U/mL penicillin, 100µg/mL streptomycin, 10mM beta-glycerophosphate, 50µg/µL ascorbic acid) for 21 days.

For adipogenic assays (n=2 mice/group), cells were seeded at 150K per well on 6-well plates in normal growth media for 24 hours prior to switching to adipogenic media (alpha-MEM with 10% FBS, 100U/mL penicillin, 100µg/mL streptomycin, 5µg/mL insulin, 0.1µM dexamethasone, 50µM indomethacin) for 7 days.

### Immunofluorescence staining and microscopy

Nesprin staining was performed on bone marrow cultures from 12-week Osx-Cre(+)/eGFP-KASH2(+) and aged-matched control mice (n=2 mice/group). After isolation and culture as described above, cells were rinsed with PBS, fixed in 4% paraformaldehyde, permeabilized with 0.3% Triton buffer and blocked with 5% goat serum (Jackson ImmunoResearch 005-000-001) prior to being tagged with a rabbit anti-nesprin2 antibody (1:300; ImmuQuest IQ565). After primary antibody tagging, cells were incubated with an anti-rabbit secondary antibody (1:300 AlexaFluor 594nm goat anti-rabbit, Invitrogen A11037), nuclei were stained with Hoechst 33342 (Invitrogen™ R37605) and visualized via fluorescent microscopy (Zeiss LSM 900). No additional staining was needed to visualize endogenous eGFP.

### Oil Red O and Alizarin Red Staining

After 7 days in adipogenic media, cultures were stained against Oil Red O as described previously [19]. Briefly, cells were rinsed with PBS, fixed in 10% neutral buffered formalin for 30 mins at room temperature, and washed with deionized water. Cultures were then incubated in 100% propylene glycol, stained with 0.5% Oil Red O in propylene glycol, and finally incubated in 85% propylene glycol. Unbound dye was removed by rinses with tap water until clear and images were taken. Oil Red O stain was then eluted with 100% propan-2-ol and optical absorbance values were read on a microplate spectrophotometer (ThermoFisher Multiskan GO) at 510nm.

After 21 days in osteogenic media, cells were rinsed in PBS, fixed with 70% ethanol, and stained with 40mM Alizarin Red solution for visualization of calcium depsosits. The stain was then eluted using 10% w/v cetylpyridium chloride (CPC) and diluted 4-fold to avoid absorbance saturation. Absorbance values were read on a microplate spectrophotometer (ThermoFisher Multiskan GO) at 562nm.

### Exercise Regimen

Eight-week old male ODLD or control mice were randomly assigned to a voluntary wheel running exercise intervention or a non-running control group without access to a running wheel (n=10/group). Animals were allotted a one-week acclimation period in these new cages followed by a six-week tracked exercise intervention. All mice were individually housed and running metrics including elapsed time, total distance, average speed, and maximum speed were measured with a cyclocomputer (CatEye Velo 7) and recorded daily. Ad libitum access to food and water was allowed. Body mass and food intake was also tracked weekly.

### Micro-computed tomography

Proximal tibiae stored frozen in PBS soaked gauze prior to scanning and were scanned immersed in PBS using an x-ray microtomograph (SkyScan 1172F). Settings as follows: power of 55kV/181mA, a voxel size of 10.33µm^3^, and 230ms integration time. The volume of interest for trabecular quantitative analysis started 10 slices distal to the proximal physis extending 1 mm. The cortical volume of interest started 2.15mm distal to the physis, extending another 0.5mm. Image reconstruction was performed with NRecon software (Bruker). After any necessary image reorientation, quantitative analysis was performed with CTAn (Bruker) and 3D image rendering was performed with CTVol.

### Biomechanical Testing

3-point bending mechanical testing was performed using an Instron Universal Testing System model 5967. Force and deflection data was collected and ran through a MATLAB code in order to determine the mechanical properties of the bone (i.e. linear flexural stiffness, maximum yield force, and maximum force).

### Statistical Analysis

Results are expressed as mean ± standard deviation. Significance was determined by 2-tailed unpaired student’s t-test or 2-way ANOVA with Tukey post-hoc test (Prism).

## Results

### Expression of EGFP-KASH2 protein disrupts LINC function in Osterix positive cells

We first crossed Osx-Cre(+) mice with the Ai14 RFP reporter line to better visualize Osx-Cre(+) animals and cells. P7 pups from this cross were visualized with an *in vivo* fluorescent scanner (IVIS® Spectrum) and notable red fluorescent in the limbs and head was visible (Figure 1A), confirming the Osx-Cre activity and specificity. We then isolated bone marrow from these animals and successfully visualized RFP containing Ai14+ cells within the bone marrow aspirates (Figure 1B), showing that we can isolate Osx+ cells with our procedure.

**Figure 1.**
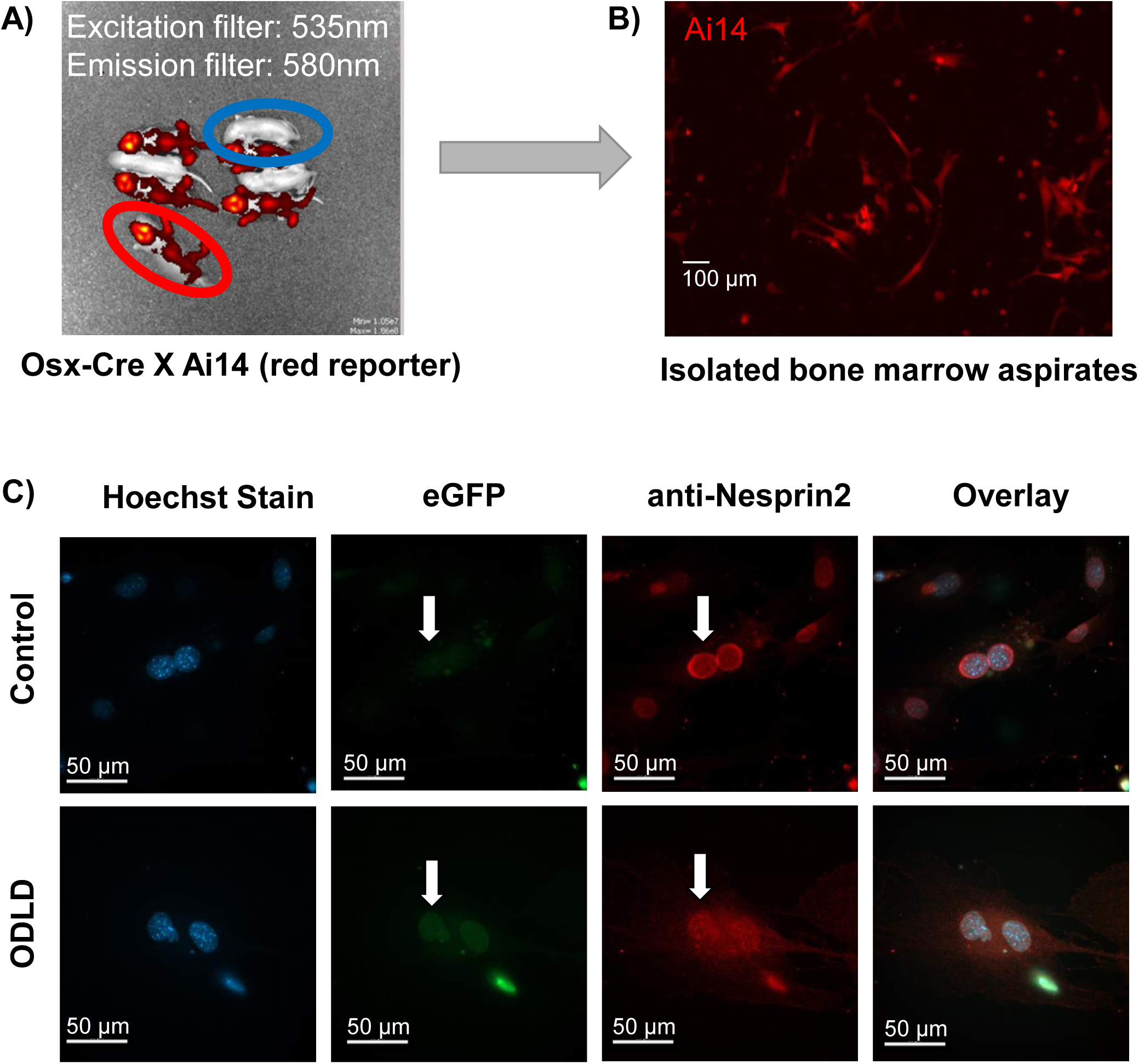
Osterix-driven LINC disruption (ODLD) model characterization. **A)** IVIS® Spectrum *in vivo* fluorescent scan of P7 pups from Osx-Cre/Ai14 cross indicating skeletal Osx-Cre activity in Osx-Cre(+) mice (red) vs. no Osx-Cre activity in Osx-Cre(-) littermates (blue). **B)** Representative image of bone marrow aspirates from 13-week male Osx-Cre(+)/Ai14 mice indicating the presence of Osx-Cre(+) cells. **C)** Bone marrow aspirates from 12-week male Osterix-driven LINC disrupted (ODLD) and control littermates. Confirmation of eGFP expression localized to the nuclear envelope with nesprin displacement compared to no eGFP visualization and intact nesprin localization to the nuclear envelope in controls.

An osterix-driven LINC disruption model was generated by crossing hemizygous Osx-Cre(+) mice with floxed Tg(CAG-LacZ/EGFP-KASH2) mice. To confirm this model works disrupts LINC function via displacing Nesprin from nuclear envelope[20], we flushed bone marrow from 12-week old animals (n= 2 mice /group) to check for endogenous eGFP expression localized to the nuclear envelope indicating the overexpression of the eGFP-KASH2 protein. However, the Cre driver for this model contains a Cre::GFP fusion protein that convolutes any interpretation of green visualization being the eGFP-KASH2 insert. Therefore, we also immunostained the cells with an anti-nesprin2 antibody to check for Nesprin-2 displacement from the nuclear envelope along with a green halo. As seen in Figure 1C, there is a clear nuclear halo of eGFP fluorescence in ODLD mice compared to no eGFP visualized in controls. Nesprin-2 staining also has a punctate and scattered appearance in ODLD mice compared to the clear nuclear positioning seen in controls indicating successful displacement of nesprins from the nuclear envelope where eGFP is present. This provides clear evidence of successful LINC disruption in cells expressing eGFP.

### Osterix-driven LINC disruption decreases osteogenic and adipogenic differentiation potential in vitro

Bone marrow cells were flushed from tibia and femur of 12-week male ODLD and negative control mice (2 animals per group, 3 technical replicates per animal), and plated for osteogenic/adipogenic differentiation assays (Figure 2A). After 7 days, adipogenic cultures were stained with Oil Red O and eluted stain was quantified via spectrophotometry at 510nm. After 21 days, osteogenic cultures were stained with Alizarin Red and eluted stain was quantified via spectrophotometry at 562nm.

**Figure 2.**
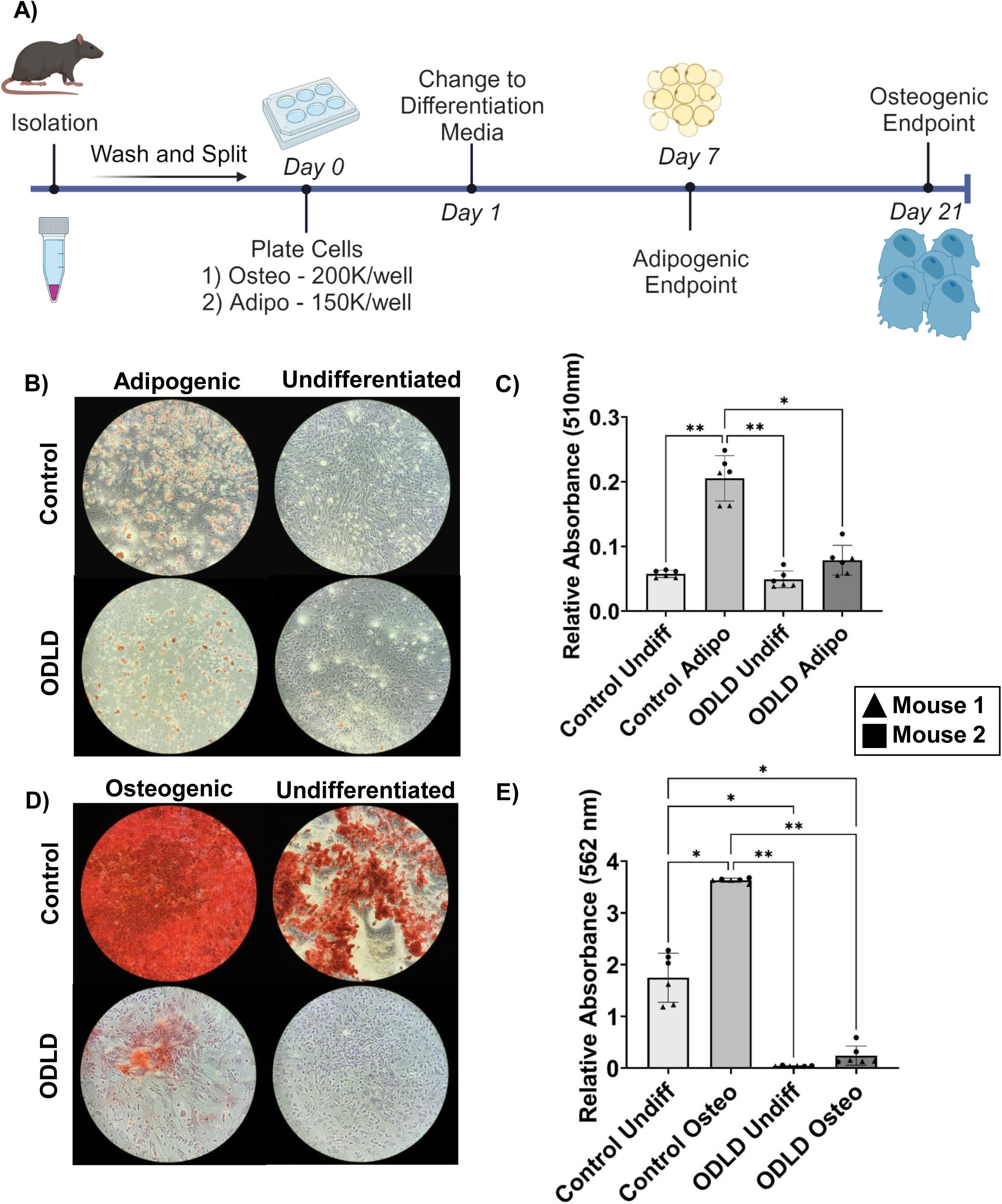
Bone marrow aspirates from ODLD mice display reduced osteogenic and adipogenic differentiation potential compared to LINC-intact controls. **A)** Schematic for differentiation assay timeline as described in Methods. **B)** Oil Red O staining after 7 days in adipogenic or normal growth media for 12 week male ODLD mice vs LINC-intact controls (n=2/grp). **C)** Absorbance values for eluted Oil Red O stain read at 510nm. n=2/group with 3 technical replicates. Shapes represent replicates from individual animals. **D)** Alizarin Red staining after 21 days in osteogenic or normal growth media. **E)** Absorbance values for eluted Alizarin Red stain read at 562nm. . n=2/group with 3 technical replicates. Shapes represent replicates from individual animals. Results presented as mean ± SD. Significance determined via ordinary one-way ANOVA. *p<0.05, **p<0.01, ***p<0.001

Shown in Fig.2B, ODLD cells showed smaller number of fat cells and relative absorbance values of Oil Red O dye from ODLD adipogenic cultures was 61.64% lower than adipogenic LINC-intact controls (p<0.05) and not significantly different from undifferentiated controls (Fig.2C). During osteogenic differentiation, ODLD cells showed markedly decreased calcium deposition as depicted by alizarin red staining (Fig.2D). Relative absorbance values of Alizarin Red stain from ODLD osteogenic cultures was 93.35% less than that of LINC-intact osteogenic controls (p<0.01), and 86.2% less than undifferentiated LINC-intact controls (p<0.05), but not significantly different from ODLD undifferentiated controls (Fig.2E). Interestingly, relative absorbance values of ODLD undifferentiated controls were 97.5% less than LINC-intact undifferentiated controls (p<0.05). Our result indicate that both the adipogenic and osteogenic differentiation potential of ODLD cells is significantly diminished compared to LINC-intact controls.

### ODLD mice trend toward decreased bone microarchitecture at 8 weeks

Prior to beginning the exercise intervention, we used an 8-week cohort of mice to establish baseline micro-CT measurements for the four possible genotypes: Osx-Cre(+)/EGFP-KASH2(+), Osx-Cre(+)/EGFP-KASH2(-), Osx-Cre(-)/EGFP-KASH2(+), Osx-Cre(-)/EGFP-KASH2(-). Animals carrying the Osx-Cre gene insert had significantly decreased bone microarchitecture compared to controls (data not shown), which is consistent with previously reported skeletal phenotypes associated with the insertion of Osx-Cre[21, 22]. Therefore, Osx-Cre(+)/EGFP-KASH2(-) mice were used as genotypic controls moving forward.

We weighed mice and harvested both tibiae and femora from Osx-Cre(+)/EGFP-KASH2(+) and Osx-Cre(+)/EGFP-KASH2(-) (n=10/group). Each tibia was cleared of soft tissue for micro-CT analysis. We noted decreased body weight in ODLD mice vs. control recorded at 17.4 ± 1.2 g and 20.7 ± 2.4 g (p=0.001), respectively, and a trend towards decreased bone microarchitecture with a 10.8% decrease in trabecular number (p<0.005), 7.13% increase in trabecular separation (p<0.05), and 8.26% decrease in total cortical cross-sectional area (p<0.05) in ODLD mice (Figure 3).

**Figure 3.**
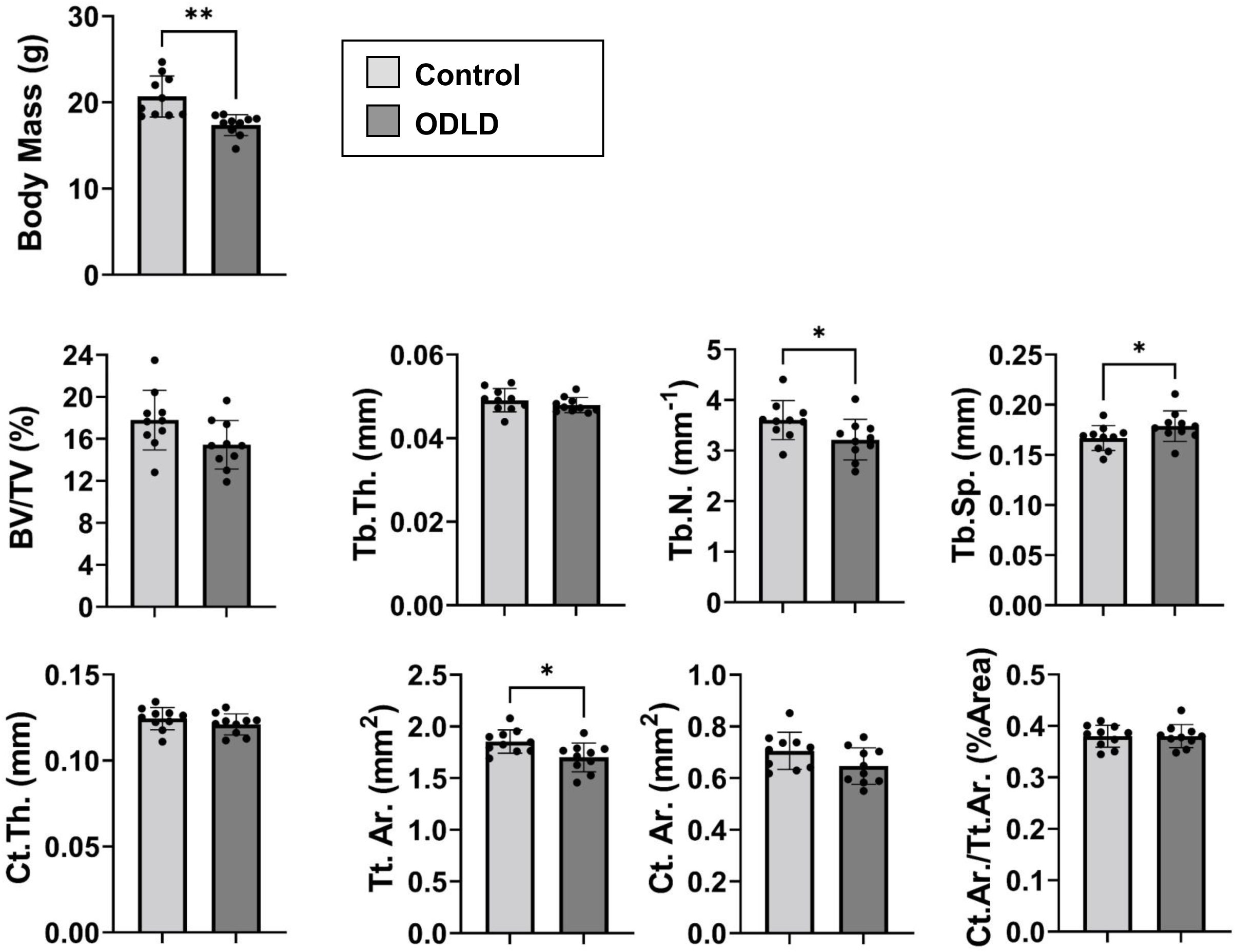
ODLD mice have less body mass and decreased trabecular number, trabecular spacing, and total cortical area of proximal tibiae in 8-week male ODLD compared to control baseline animals. Trabecular measurements include bone volume fraction (BV/TV), trabecular thickness (Tb.Th.), trabecular spacing (Tb.Sp.), and trabecular number (Tb.N). Cortical measurements include total cross-sectional area inside the periosteal envelope (Tt.Ar.), cortical bone area (Ct.Ar.), cortical area fraction (Ct.Ar./Tt.Ar.), and average cortical thickness (Ct.Th.) Significance determined via non-parametric Student’s t-test. Decreased body weight in ODLD mice vs control recorded at 17.4 ± 1.2 g and 20.7 ± 2.4 g, respectively (p=0.001) and a trend towards decreased bone microarchitecture with a 10.8% decrease in trabecular number (p<0.005), 7.13% increase in trabecular separation (p<0.05), and 8.26% decrease in total cortical cross-sectional area (p<0.05) in ODLD mice. *p<0.05, **p<0.01, ***p<0.001

### Osterix-driven LINC disrupted mice run longer, farther, and faster than controls

8-week male mice were placed in a cage equipped with a running wheel (or without) and given a one-week acclimation period before beginning the six-week tracked exercise intervention (Fig.4A). No changes in body mass and food intake were detected during the running protocol (Fig.4B). Tracked running metrics including elapsed time, average distance, average velocity, and max speed were recorded daily. As shown in Fig.4D, our data indicate that ODLD mice ran significantly longer (170.4 ± 15.0 mins vs 133.7 ± 9.2 mins, p<0.001), farther (2.041 ± 0.3 km vs to 1.62 ± 0.26 km, p<0.05), and faster (47 ± 0.04 km/hr vs 0.41 ± 0.02 km/hr, p<0.01) when compared to LINC-intact counterparts.

**Figure 4.**
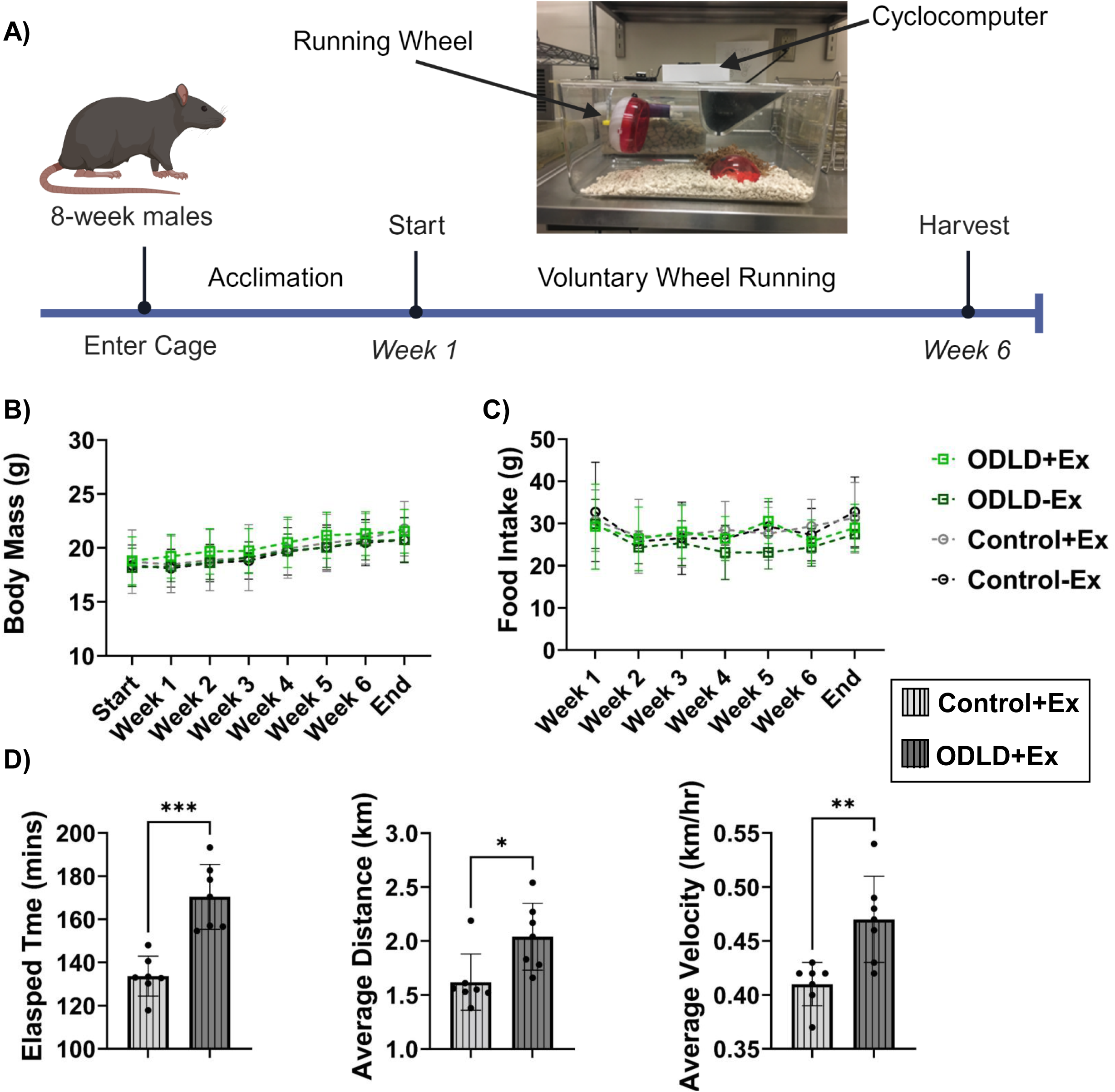
ODLD mice run significantly longer, farther, and faster than controls. **A)** Schematic for voluntary wheel running intervention timeline with running cage set-up **B)** Body mass steadily increased throughout exercise intervention with no significant difference between groups. **B)** No significant difference in food intake between groups. **C)** ODLD mice ran longer, farther, and faster than LINC-intact controls. Running metrics tracked by cyclometer measuring elapsed time (mins/day), average distance (km/day), average velocity (km/hr/day), and max velocity. Significance determined via 2-way ANOVA for body weight and food intake and unpaired Student’s t-test for elapsed time, average distance, and average velocity. *p<0.05, **p<0.01, ***p<0.001

### LINC disruption increases trabecular thickness but does not affect additional bone microarchitecture or biomechanical properties in mice subjected to six-week running protocol

Shown in Fig.5A, bone microarchitecture was measured via micro-CT to assess any effects of LINC disruption on exercise-induced bone accrual in the proximal tibial metaphysis and diaphysis for trabecular and cortical parameters, respectively. There was no notable difference in trabecular bone fraction (BV/TV), trabecular separation (Tb.Sp.), trabecular number (Tb.N), total cortical tissue area (Tt.Ar.), cross-sectional area (Ct.Ar.), cortical area fraction (Ct.Ar./Tt.Ar.), or cortical thickness (Ct.Th.), but there was a 4.2% (p<0.05) increase in trabecular thickness (Tb.Th.) in exercised ODLD mice compared to sedentary ODLD controls. Further, no changes were found during biomechanical tests (Fig.5B).

**Figure 5.**
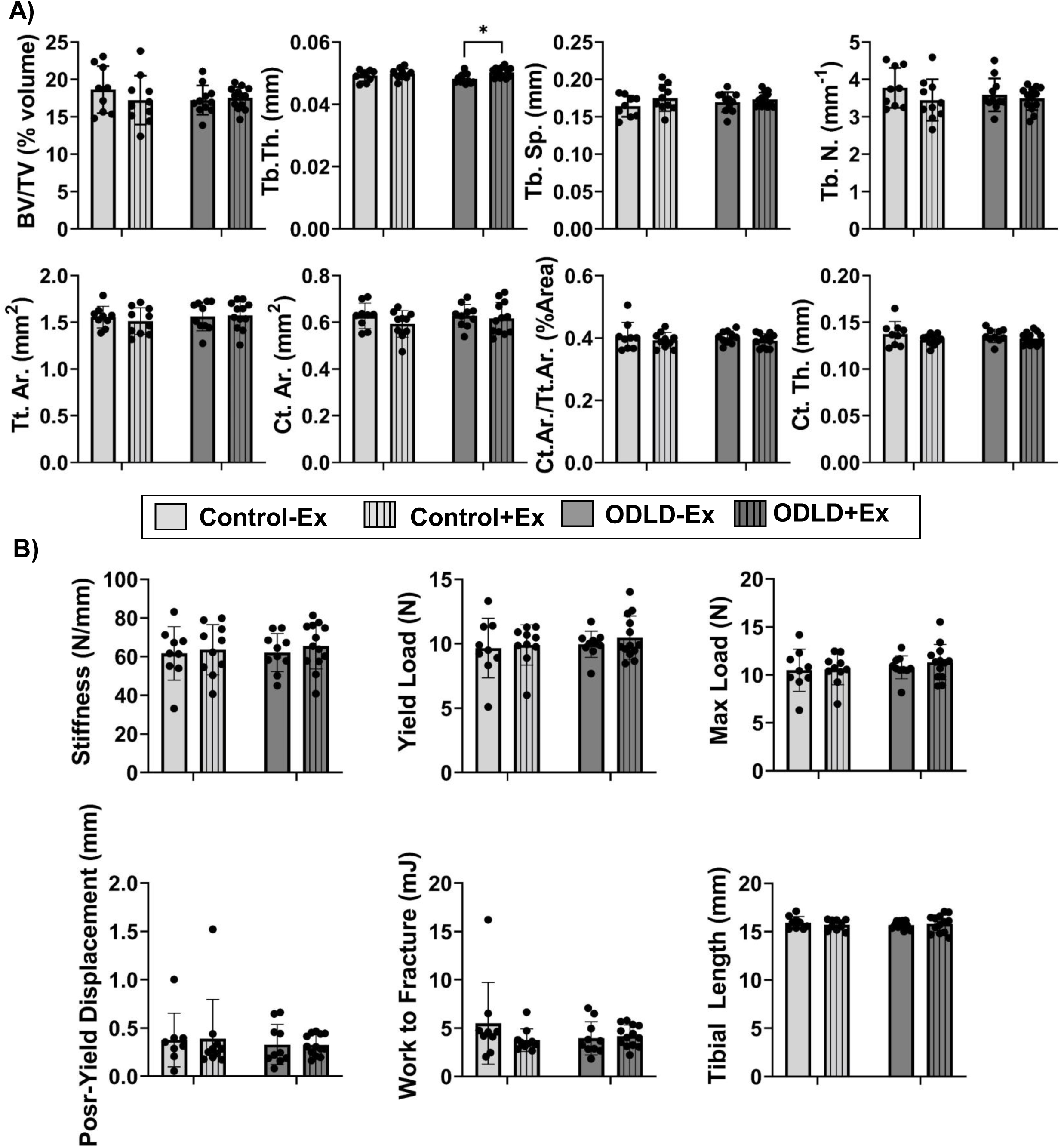
Trabecular and cortical microarchitecture of proximal tibiae were unaffected by LINC disruption after 6-week voluntary wheel running. **A)** Trabecular measurements include bone volume fraction (BV/TV), trabecular thickness (Tb.Th.), trabecular spacing (Tb.Sp.), and trabecular number (Tb.N). Cortical measurements include total cross-sectional area inside the periosteal envelope (Tt.Ar.), cortical bone area (Ct.Ar.), cortical area fraction (Ct.Ar./Tt.Ar.), and average cortical thickness (Ct.Th.) **B**) Tibial biomechanical properties and length post-exercise. *p<0.05, **p<0.01, ***p<0.001

## Discussion

The role of LINC complex in mechanical signaling of bone progenitor cells has been well studied *in vitro*, but its function *in vivo* remains underexplored. Here we sought to elucidate if LINC complex disruption in *Osx*(+) pre-osteoblastic cells compromises cell phenotype, bone quality and response to voluntary wheel running. After successfully generating a LINC disruption mouse model, we found decreased osteogenic and adipogenic differentiation potential in ODLD cells as well as diminished trabecular architecture in ODLD mice at an 8-week baseline. There were no remarkable differences in bone microarchitecture or biomechanical properties of exercised and sedentary, non-exercised mice for both ODLD and LINC-intact controls.

Similar studies using a six week voluntary wheel running intervention but using older (16 week) female mice with diet-induced obesity or caloric restriction found increased trabecular quality and quantity post-exercise in animals on a regular diet [23] [24]. While bones of male mice also shown to respond to voluntary wheel running [25] [26] [27], other studies using male mice reported minimal effects of exercise on bone after 10 or 21 weeks of voluntary wheel running [28, 29]. Compared to reported daily running metrics in these studies (ranging from 4-10 km/day), our mice ran less on average (1.8 km/day). While we do not know why our mice run less, this may have affected our outcomes. Furthermore, our mice were 8-weeks old when placed in the exercise intervention, which is before the 12-week reported age of skeletal maturity [28, 30]. Therefore, it is possible that mechanical challenge provided by voluntary wheel running did not affect the ongoing bone growth in these mice. We found a slight increase in trabecular thickness (Tb.Th.) in ODLD mice that ran 26% percent more than controls suggesting a more robust running protocol may be needed in future studies to see possible effects.

Disruption of LINC complex via KASH overexpression *in vitro* decreases osteogenic activity in human bone marrow mesenchymal stem cells by reducing Runx2 via increased histone acetylation [32]. Our data further indicates that KASH mediated LINC disruption also decreases adipogenic potential *in vitro,* suggesting that LINC may play a role in cell fate determination. Our recent findings indicate that both disruption Lamin A/C[33] and LINC function via co-depletion of SUN1/2 decreases adipogenesis, in the case of the latter this happens via accumulation of H3K9me3 at the adipogenic loci [34]. While we had a strong effect *in vitro*, cell microenvironment on plastic does not adequately mimic the bone marrow microenvironment; therefore, *in vitro* findings may not adequately reflect the *in vivo* situation. This combined with i*n vivo* observation of decreased bone indices at 8-weeks suggests that LINC disruption may affect cell differentiation in pre-osteoblasts early on but not at later stages, as we did not see any effect in sedentary animals at 15 weeks. Further studies using different modalities of LINC disruption (i.e. SUN & Nesprin depletion models) or the use of different Cre drivers (i.e. Prrx1, Col1a1, Ocn, etc.) would shed further light on the impact of impaired mechanotransduction in skeletal stem/progenitor cells.

## Acknowledgements

This study was supported by NIH AG059923, AR075803, P20GM109095 and NSF 2025505. The author(s) declare no competing interests financial or otherwise.

